# Epitope - based peptide vaccine against glycoprotein GPC precursor of *Lujo virus* using immunoinformatics approaches

**DOI:** 10.1101/2020.09.09.287771

**Authors:** Arwa A. Mohammed, Mayada E. Elkhalifa, Khadija E. Elamin, Rawan A. Mohammed, Musab E. Ibrahim, Amina I. Dirar, Sara H. Migdar, Maha A. Hamid, Emeirii H. Elawad, Salam O. Abdelsalam, Mohamed A. Hassan

**Affiliations:** Department of Biotechnology, Africa city of Technology, Khartoum, Sudan; Department of Pharmacy, Sudan Medical Council, Khartoum, Sudan; Department of Pharmaceutics, Faculty of Pharmacy, Alneelain University, Khartoum, Sudan; Department of Botany, Faculty of science, University of Khartoum, Sudan; Department of Bioinformatics, Faculty of Information & Science Technology, Multimedia University, Malaysia; Department of Clinical and industrial pharmacy, Faculty of Pharmacy, National university, Khartoum, Sudan; Department of Pharmaceutical Chemistry, Faculty of Pharmacy, University of Khartoum, Sudan; Department of Clinical Pharmacy, School of Pharmacy, Ahfad University for Women, Khartoum, Sudan; Department of Pharmacology, Faculty of Medicine, University of Khartoum, Sudan; Department of Clinical Immunology, Sudan Medical Specialization Board, Khartoum, Sudan; Department of Bioinformatics, DETAGEN Genetics Diagnostic Center, Kayseri, Turkey

**Keywords:** Immunoinformatics, glycoprotein GPC precursor, Epitope-based vaccine, *Lujo virus* LUJV, VHF

## Abstract

**Background:** *Lujo virus* (LUJV) is a highly fatal human pathogen belonging to the *Arenaviridae* family. *Lujo virus* causes viral hemorrhagic fever (VHF). An *In silico* molecular docking was performed on the GPC domain of *Lujo virus* in complex with the first CUB domain of neuropilin-2.

**The aim** of this study is to predict effective epitope-based vaccine against glycoprotein GPC precursor of *Lujo virus* using immunoinformatics approaches.

**Methods and Materials:** glycoprotein GPC precursor of *Lujo virus* Sequence was retrieved from NCBI. Different prediction tools were used to analyze the nominee’s epitopes in BepiPred-2.0: Sequential B-Cell Epitope Predictor for B-cell, T-cell MHC class II & I. Then the proposed peptides were docked using Autodock 4.0 software program.

**Results and Conclusions:** The proposed and promising peptides FWYLNHTKL and YMFSVTLCI shows a very strong binding affinity to MHC class I & II alleles with high population coverage for the world, South Africa and Sudan. This indicates a strong potential to formulate a new vaccine, especially with the peptide YMFSVTLCI which is likely to be the first proposed epitope-based vaccine against glycoprotein GPC of *Lujo virus*. This study recommends an in-vivo assessment for the most promising peptides especially FWYLNHTKL, YMFSVTLCI and LPCPKPHRLR.

## 1. Introduction

Arenaviruses are rodent-borne viruses. A genetically unique arenavirus called *Lujo virus*, has been discovered as the causal agent of a nosocomial outbreak of acute febrile disease with hemorrhagic manifestations in Zambia and South Africa. The outbreak had a high case fatality rate of almost 80% ^[1]^. These viruses are genetically and geographically divided to the Old World mammarena viruses, endemic to West Africa, and the New World mammarena viruses, endemic to South and North America ^[2]^.

*Lujo virus* causes viral hemorrhagic fever (VHF) which can be caused by five distinct families of viruses: the filo-, arena-, flavi-,rhabdo- and bunya virus family ^[3]^.

Viral hemorrhagic fever (VHF) is an acute systemic illness classically involving fever, a constellation of initially nonspecific signs and symptoms, and a propensity for bleeding and shock. With *Lujo virus,* hemorrhagic fever (LVHF) illness typically began with the abrupt onset of fever, malaise, headache, and myalgias followed successively by sore throat, chest pain, gastrointestinal symptoms, rash, minor hemorrhage, subconjunctival injection, and neck and facial swelling over the first week of illness. No major hemorrhage was noted. Neurological signs were sometimes seen in the late stages. Shock and multi-organ system failure, often with evidence of disseminated intravascular coagulopathy, ensued in the second week, with death in four of the five cases ^[4]^. There are currently limited preventative and therapeutic options for patients infected with these highly pathogenic viruses ^[5]^.

Arenaviruses are enveloped negative-strand RNA viruses with a genome that is bi-segmented into S and L segments. The S segment encodes a nucleocapsid protein (NP) and an envelope glycoprotein precursor (GPC); the L segment encodes a matrix protein (Z) and an RNA-dependent RNA polymerase (L). The GPC is synthesized as a single polypeptide and undergoes processing by the host cell signal peptidase (SPase) and subtilisin-like kexinisozyme-1/site-1-protease (SKI-1/S1P), yielding typical receptor binding (G1), transmembrane fusion (G2), and stable signal peptide (SSP) subunits, respectively ^[6–8]^. Viral entry into target cells is initiated by the binding of G1 to appropriate cell surface receptors. The first cellular receptor for arenavirus to be identified was_-dystroglycan (_DG), a ubiquitous receptor for extracellular matrix proteins ^[9]^.

The Understanding of epitope/antibody interaction is the key to constructing potent vaccines and effective diagnostics. The host defense mechanisms against viruses generally vary from germline-encoded immunity, present early in the evolution of microorganisms to activate and induce specific adaptive immune responses by the production of Th-1 andTh-2 cytokines. B-cells recognize antigens via membrane bound antibodies using B-cell receptors (BCRs), resulting in the secretion of antibodies that bind to the antigen and deactivate or remove it. Processing and presentation of peptide epitopes are essential steps in cell-mediated immunity ^[10]^. *Lujo virus* (LUJV) is a highly fatal human pathogen belonging to the *Arenaviridae* family. This virus is unique; as it uses neuropilin-2 (NRP2) as a cellular receptor. Previous study revealed that the GP1 receptor-binding domain of LUJV (LUJVGP1) recognizes NRP2, where its recognition is metal-ion dependent. The binding of a Ca2^+^ ion stabilizes the conformations of Asp127 and Glu79 from NRP2, pre-organizing them for interaction with Lys110 of LUJVGP1. CUB domain of NRP2 is almost completely conserved among humans, mice, rats and bats, and the only slight variations occur outside of the binding site for LUJV. Hence all of these animal species have a potential to serve as reservoirs for LUJV, considering only the compatibility to NRP2 ^[2]^. *In silico* molecular docking was performed on the GP1 domain of *Lujo virus* in complex with the first CUB domain of neuropilin-2 ^[2]^.

The aim of the study is to predict effective epitope-based vaccine against envelope glycoprotein precursor (GPC); of *Lujo virus*. The development of immunogenetics will enhance apprehension of the impact of genetic factors on the interindividual and interpopulation variations in immune responses to vaccines that could be helpful to progress new vaccine strategies ^[11]^. In silico/reverse vaccinology had replaced conventional culture-based vaccines because it reduces the cost required for laboratory investigations of pathogens and speeds up the time needed to achieve the results ^[12,13]^.

Therefore, using immunoinformatics approaches to predict this new kind of vaccines could provide magnificently valuable insights needed for the prevention of *Lujo virus*. Normally, the investigation of the binding affinity of antigenic peptides to the MHC molecules is the main goal when predicting epitopes. Using such tools and information leads to the development of new vaccines. While these approaches permit the optimization of a vaccine for a specific population, the problem can also be reformulated to design a “universal vaccine”, one that provides maximum coverage for the whole worlds’ population ^[14,15]^. In this study, we focused on both MHC class II and class I along with carrying out molecular docking in HLA-A0201.

## 2. Materials and Methods

### 2.1 Sequences retrieval

The amino acids sequences of Glycoprotein GPC (Glycoside hydrolase family) of *Lujo virus* were retrieved from NCBI database (https://www.ncbi.nlm.nih.gov/protein) ^[16]^ in FASTA format on July 2018. Different prediction tools of Immune Epitope Database IEDB analysis resource (http://www.iedb.org/) ^[17]^ were then used to analyze the candidate epitopes as shown in figure (1).

**Figure 1:**
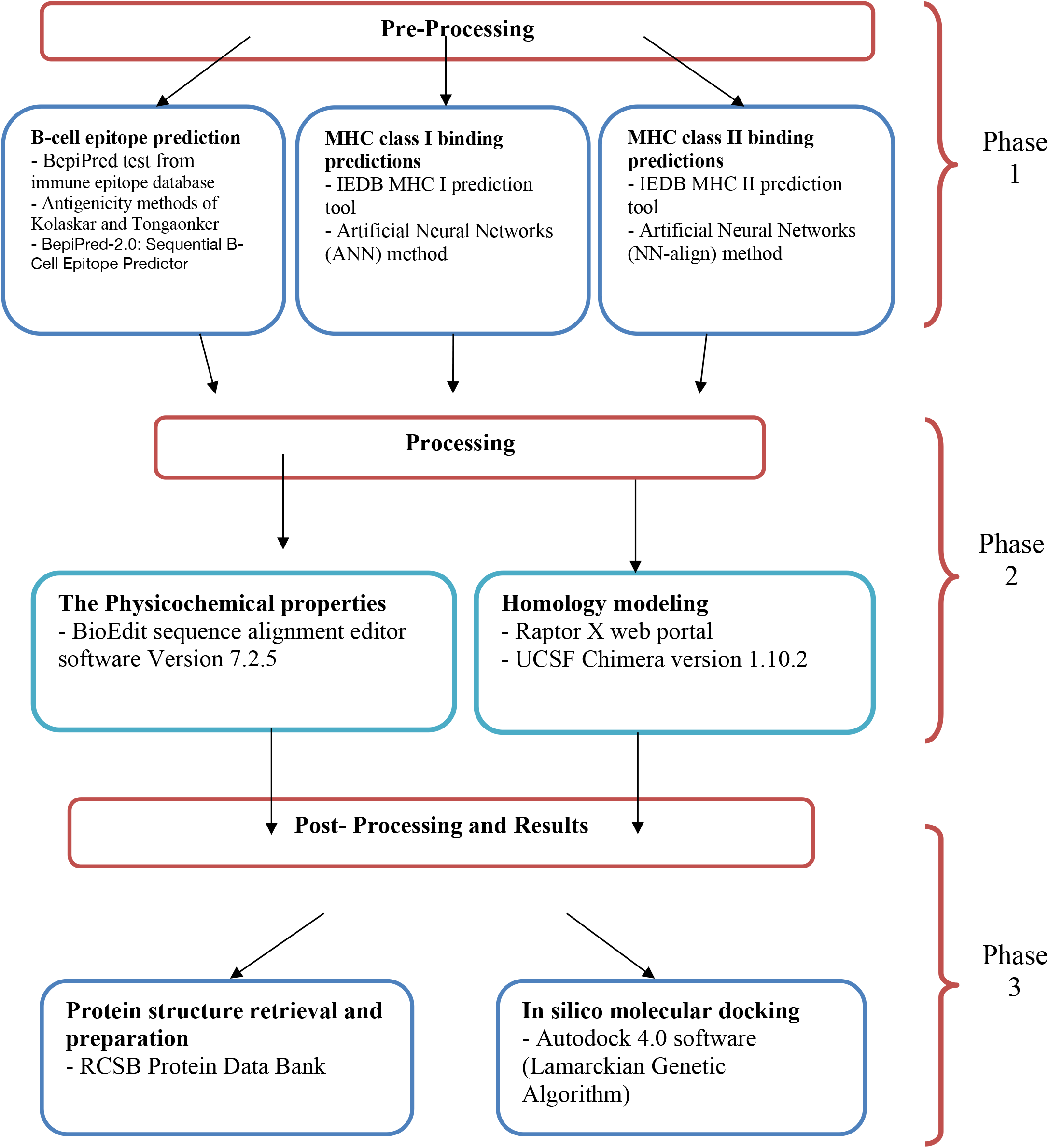
The three phases of Material and Method process.

### 2.2 Conservation region and physicochemical properties

Conservation regions were determined using multiple sequence alignment with the help of Clustal-W in the Bioedit software version ^[18]^. Epitope conservancy prediction for individual epitopes was then calculated using the IEDB analysis resource. Conservancy can be defined as the portion of protein sequences that restrain the epi-tope measured at or exceeding a specific level of identity ^[19]^. The physicochemical properties of the retrieved sequence; molecular weight and amino acid composition; were also determined by Bioedit software version.

### 2.3 B cell epitope prediction tools

Candidate epitopes were analyzed using several B-cell prediction methods to determine the antigenicity, flexibility, hydrophilicity and surface accessibility. The linear prediction epitopes were obtained from Immune epitope database (http://tools.iedb.org/bcell/result/) ^[20]^ by using BepiPred test with a threshold value of 0.149 and a window size 6.0

Moreover, surface accessible epitopes were predicated with a threshold value of 1.0 and window size 6.0 using the Emini surface accessibility prediction tool ^[21]^.

Kolaskar and Tongaonker antigenicity methods (http://tools.iedb.org/bcell/result/) were proposed to determine the sites of antigenic epitopes with a default threshold value of 1.030 and a window size 6.0 ^[22]^.

### 2.4 T cell epitope prediction tools

#### 2.4.1 Peptide binding to MHC class I molecules

The peptide binding was assessed by IEDB MHC class I prediction tool at http://tools.iedb.org/mhc1. This tool employs different methods to determine the ability of submitted sequence to bind to a specific MHC class I molecule. The artificial neural network (ANN) method ^[23, 24]^ was used to calculate IC50 values of peptide binding to MHC-class I molecules. For both frequent and non-frequent alleles, peptide length was set to 9 amino acids earlier to the prediction. The alleles having binding affinity IC50 equal to or less than 500 nM were considered for further analysis.

#### 2.4.2 Peptide binding to MHC class II molecules

To predict the peptide binding to MHC class II molecules, MHC class II prediction tool http://tools.iedb.org/mhcII provided by Immune Epitope Database (IEDB) analysis resource and human allele references set was used. The Artificial Neural Network prediction method was chosen to identify the binding affinity to MHC class II grooves and MHC class II binding core epitopes. All epitopes that bind to many alleles at score equal to or less than 1000 half-maximal inhibitory concentration (IC50) were selected for further analysis.

### 2.5 Population coverage

Population coverage for each epitope was calculated by the IEDB population coverage tool at http://tools.iedb.org/tools/population/iedb_input ^[25]^. This tool is aimed to determine the fraction of individuals predicted to respond to a given set of epitopes with known MHC restrictions. For every single population coverage, the tool computed the following information: (1) predicted population coverage, (2) HLA combinations recognized by the population, and (3) HLA combinations recognized by 90% of the population (PC90). All epitopes and their MHC class I and MHC class II molecules were assessed against population coverage area selected before submission.

### 2.6 Homology modeling

The 3D structure of glycoprotein GPC of *Lujo virus* was predicted using Raptor X web portal (http://raptorx.uchicago.edu/) ^[26]^. The reference sequence was submitted in FASTA format on 14/9/2018 and structure received on 15/9/2018. Structure was then treated with UCSF Chimera 1.10.2 ^[27]^ to visualize the position of proposed peptides.

### 2.7 In silico Molecular Docking

#### 2.7.1 Ligand Preparation

In order to estimate the binding affinities between the epitopes and molecular structure of MHC class I & MHC class II, in silico molecular docking was used. Sequences of the proposed epitopes were selected from *Lujo virus* reference sequence using Chimera 1.10 and saved as pdb file. The obtained files were then optimized and the energy was minimized. The HLA-A0201 was selected as the macromolecule for docking. Its crystal structure (4UQ3) was downloaded from the RCSB Protein Data Bank (http://www.rcsb.org/pdb/home/home.do), which was in complex with an azobenzene-containing peptide ^[28]^.

The crystal structure of LUJVGP1/NRP2 was retrieved from protein databank (PDB ID: 6GH8) ^[2]^. Molecular docking was performed using Autodock 4.0 software, based on Lamarckian Genetic Algorithm; which combines energy evaluation through grids of affinity potential to find the suitable binding position for a ligand on a given protein ^[29]^. Polar hydrogen atoms were added to the protein targets and Kollman united atomic charges were computed. All hydrogen atoms were added to the ligands before the Gastiger partial charges were assigned. The co-crystal ligand was removed and the bond orders were checked. The target’s grid map was calculated and set to 60×60×60 points with grid spacing of 0.375 Ǻ. The grid box was then allocated properly in the target to include the active residue in the center. The default docking algorithms were set in accordance with standard docking protocol ^[30]^. Finally, ten independent docking runs were carried out for each ligand and results were retrieved as binding energies. Poses that showed lowest binding energies were visualized using UCSF chimera ^[31]^.

## 3. Results

### 3.1. *Lujo virus* glycoprotein GPC physical and chemical parameters

The physicochemical properties of the *Lujo virus* glycoprotein GPC protein was assessed using BioEdit software version 7.0.9.0. The protein length was found to be 454 amino acids. The amino acids that form *Lujo virus* glycoprotein GPC protein is shown in Figure (2) along with their numbers and molar percentages in (Mol%).

**Figure 2:**
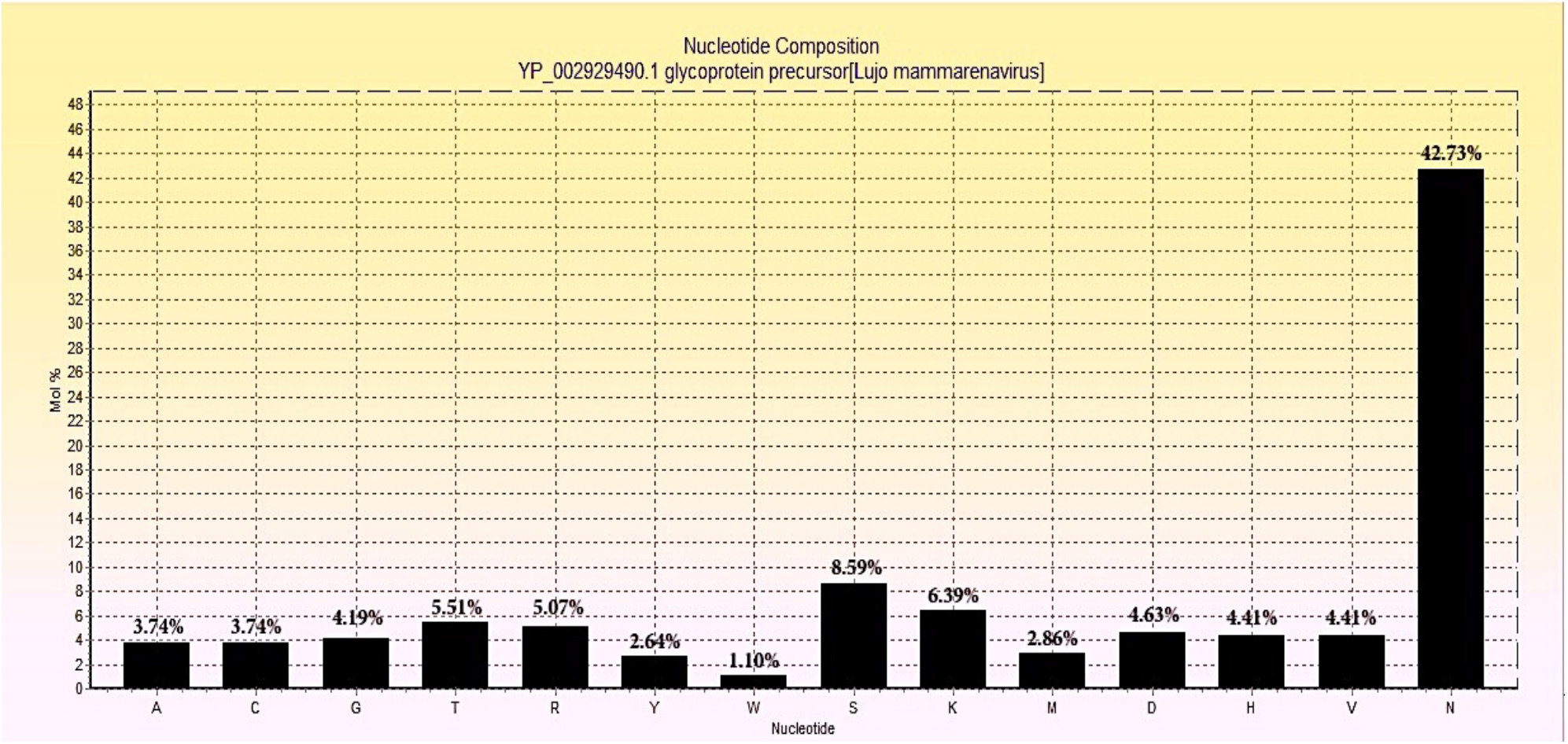
Amino acids composition of *Lujo virus* glycoprotein GPC using Bioedit

### 3.2 B-cell epitope prediction

The ref sequence of the *Lujo virus* glycoprotein GPC was subjected to a Bepipred linear epitope prediction. Emini surface accessibility, Kolaskar and Tongaonkar antigenicity methods in IEDB were used to determine bindings to the B cell, testing its surface and immunogenicity. The results are shown in Table1.

**Table 1.**
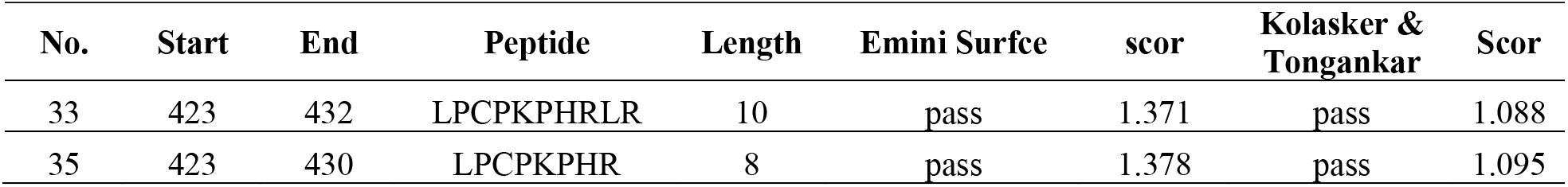
Most linear epitopes, high surface accessibility and immunogenicity bindings to the B cell.

### 3.3. Prediction of T helper cell epitopes and interaction with MHC class I alleles

*Lujo virus* glycoprotein GPC sequence was analyzed using IEDB MHC class I binding prediction tool based on ANN-align with half-maximal inhibitory concentration (IC_50_) ≤**500**; the least most promising epitopes that had a binding affinity with the Class I alleles along with their positions in the *Lujo virus* glycoprotein GPC are shown in Table 2.

**Table 2.**
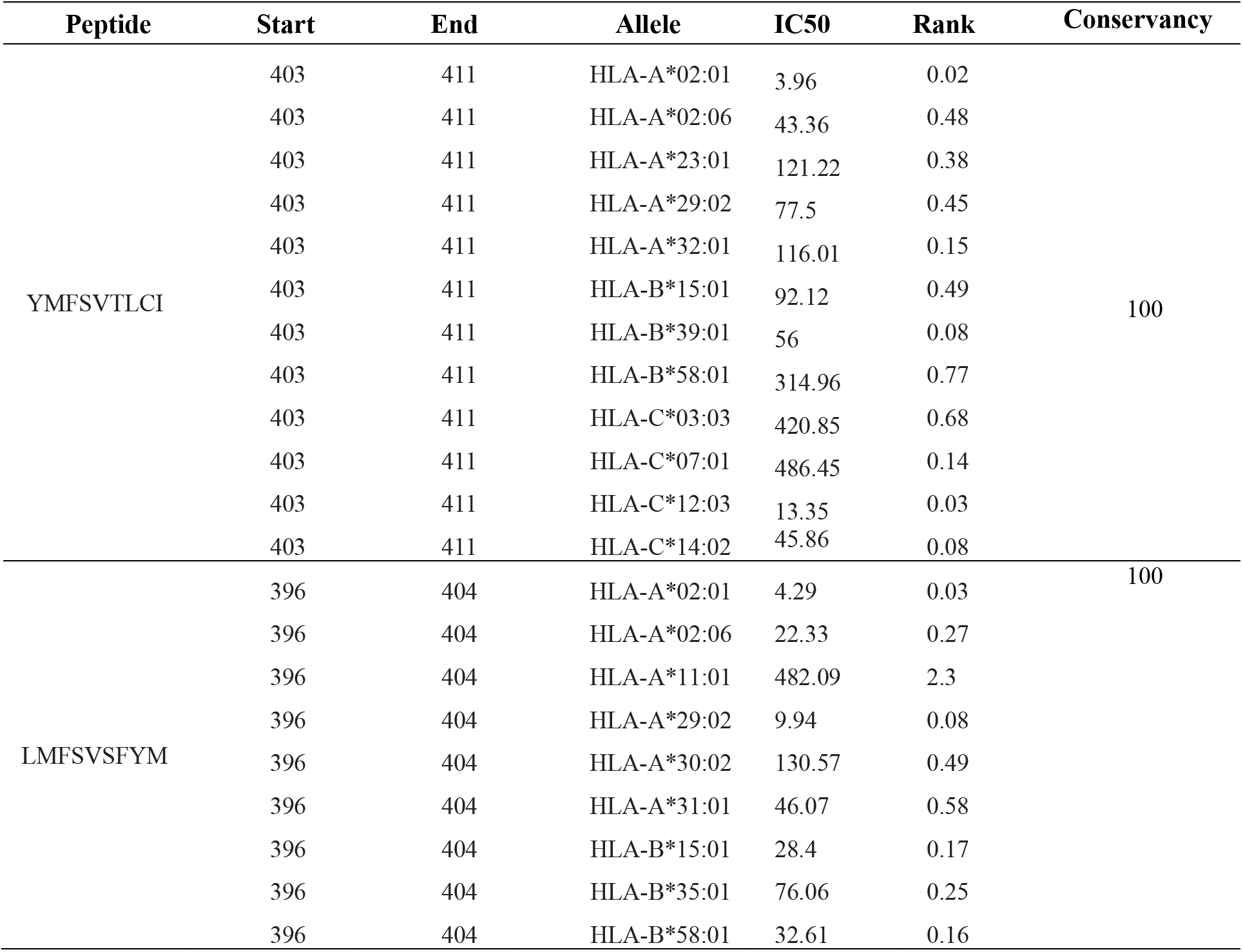
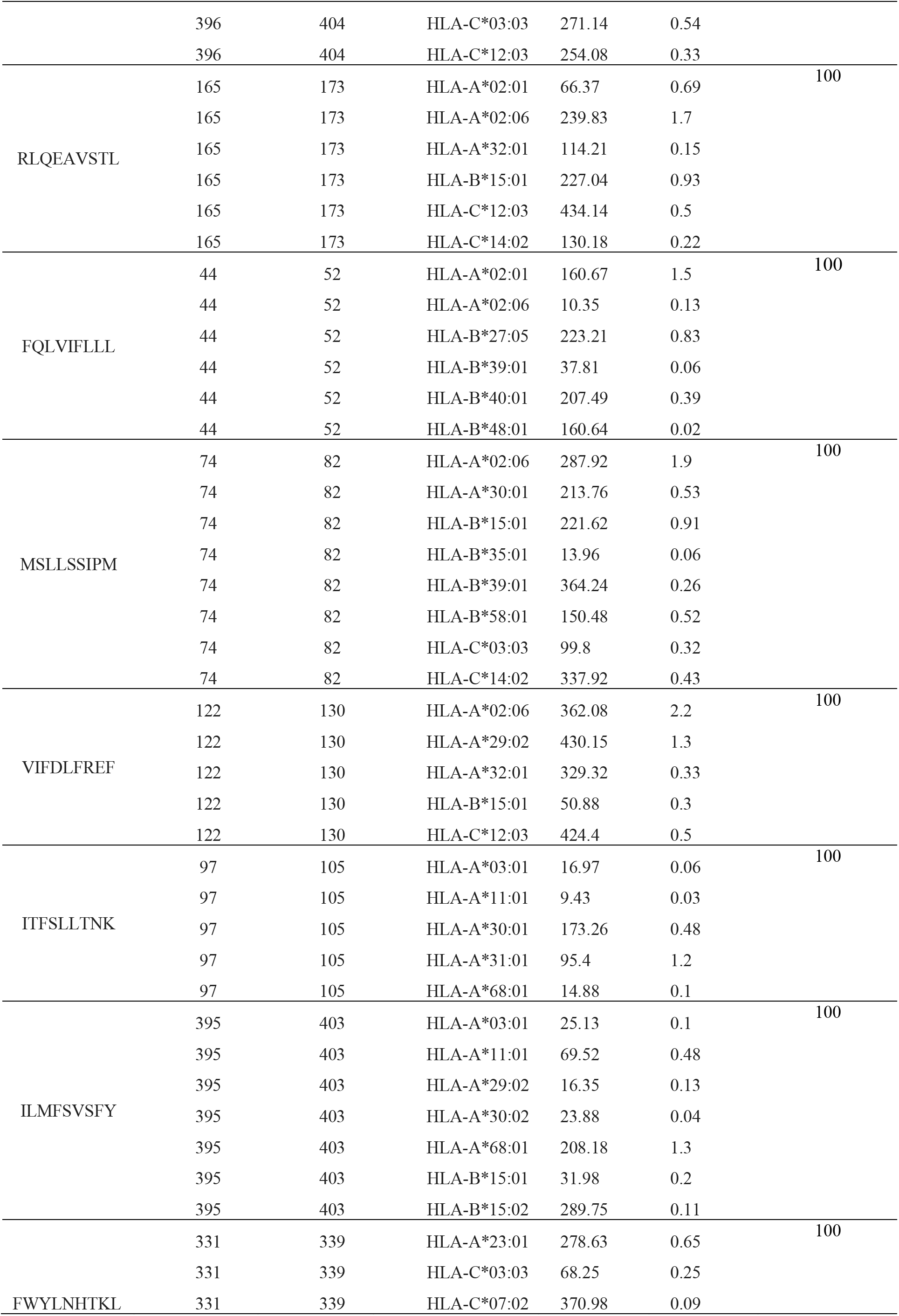
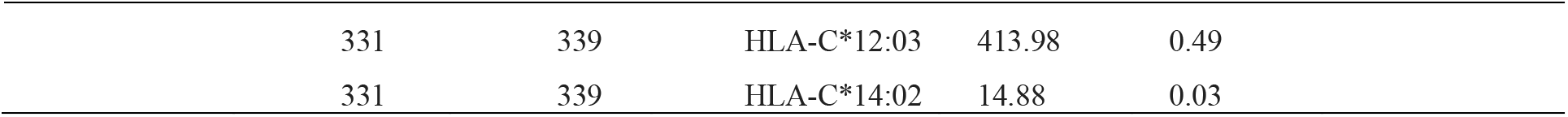
Most potential T-cell epitopes with interacting MHC-class I alleles, their positions, IC50, rank and conservancy

### 3.4. Prediction of T helper cell epitopes and interaction with MHC class II alleles

*Lujo virus* glycoprotein GPC sequence was analyzed using IEDB MHC class II binding prediction tool based on NN-align with half-maximal inhibitory concentration (IC_50_) ≤**1000**. The list of the epitopes and their correspondent bindings to MHC class II alleles, along with their positions in the *Lujo virus* glycoprotein GPC. While the list of the most promising epitopes, that had a strong binding affinity to MHC class II alleles and depending on the number of their binding alleles are shown in Table 3.

**Table 3.**
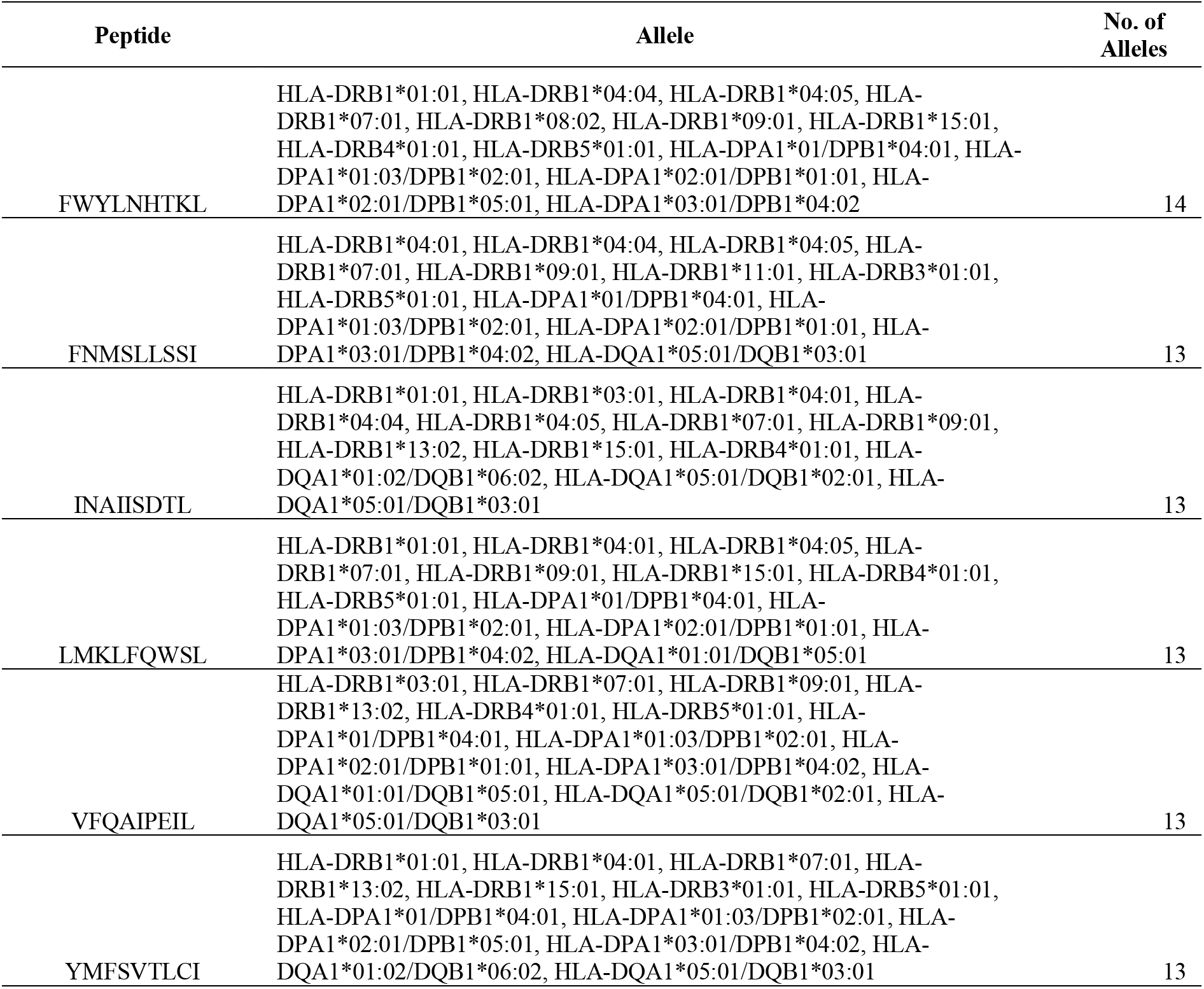
Most potential T-cell epitopes with interacting MHC-class II alleles

### 3.5 Population coverage

A population coverage test was performed to detect all the epitopes that bind to MHC class I alleles and MHC class II alleles available in the database in relation to the world, South **Africa**, and Sudan.

### 3.6. 3D structure

### 3.7. Molecular Docking

Three peptides; FWYLNHTKL, LPCPKPHRLR and YMFSVTLCI were docked onto protein target of GP1 domain of *Lujo virus* in complex with the first CUB domain of NRP2.

## 4. Discussion

In this computational immunoinformatic study we suggest a new promising highly selective peptide vaccine against *Lujo virus* for the first time according to our knowledge. We expect to obtain a peptide-based vaccine with high antigenicity and minimum allergic effect rather than the currently used vaccines. This challenge starts with having good information about the protein structure of *Lujo virus* from literature review, then the reference sequence of *Lujo virus* glycoprotein GPC was obtained from the NCBI. To determine the binding affinity of the conserves epitopes to B-cell and to examine the immunogenicity we subjected the reference sequence of *Lujo virus* glycoprotein GPC to IEDB database. Bepipred linear epitope prediction test, Emini surface accessibility test and Kolaskar and Tongaonkar antigenicity test were examined. For Bepipred test of B-cell the total number of epitopes was 39. For Emini surface accessibility prediction, 29 conserved epitopes were passing the default threshold 1.0. In Kolaskar and Tongaonkar antigenicity, 7 epitopes gave scores above the default threshold 1.045. However, there are only two epitopes that pass the three tests (LPCPKPHRLR, LPCPKPHR). The reference glycoprotein GPC strain was analyzed using IEDB class MHC-class I binding prediction tool to predict T cell epitope. 165 peptides were predicted to interact with different class MHC-class I alleles. For class MHC-class II binding prediction there were 315 epitopes found to interact with class MHC-class II alleles. The peptides YMFSVTLCI, LMFSVSFYM, RLQEAVSTL, FQLVIFLLL, MSLLSSIPM, VIFDLFREF, ITFSLLTNK, ILMFSVSFY and FWYLNHTKL had the affinity to bind the highest number of class MHC-class I alleles. The peptides FWYLNHTKL, YMFSVTLCI, FNMSLLSSI, INAIISDTL, LMKLFQWSL and VFQAIPEIL had the affinity to bind the highest number of class MHC-class II alleles. The most promising three peptides for both class MHC-class I and MHC-class II were FWYLNHTKL, LPCPKPHRLR and YMFSVTLCI as shown in figure (3). On the other hand, the world population coverage of all epitopes that bind to MHC-class I was found to be 99.83%, while the world population coverage of all epitopes that bind to MHC-class II were 68.23% as presented in table 4. For the binding affinity to MHC-class I and MHC-class II the peptide FWYLNHTKL was found to bind 14 different alleles of MHC-class II & five alleles of MHC-class I, that gave a world population coverage of 74.82%, 43.01% for South Africa and 51.56% for Sudan for both MHC class I and II as shown in table 5. This finding shows a very strong potential to formulate an epitope-based peptide vaccine for *Lujo virus*.

**Table 4.**
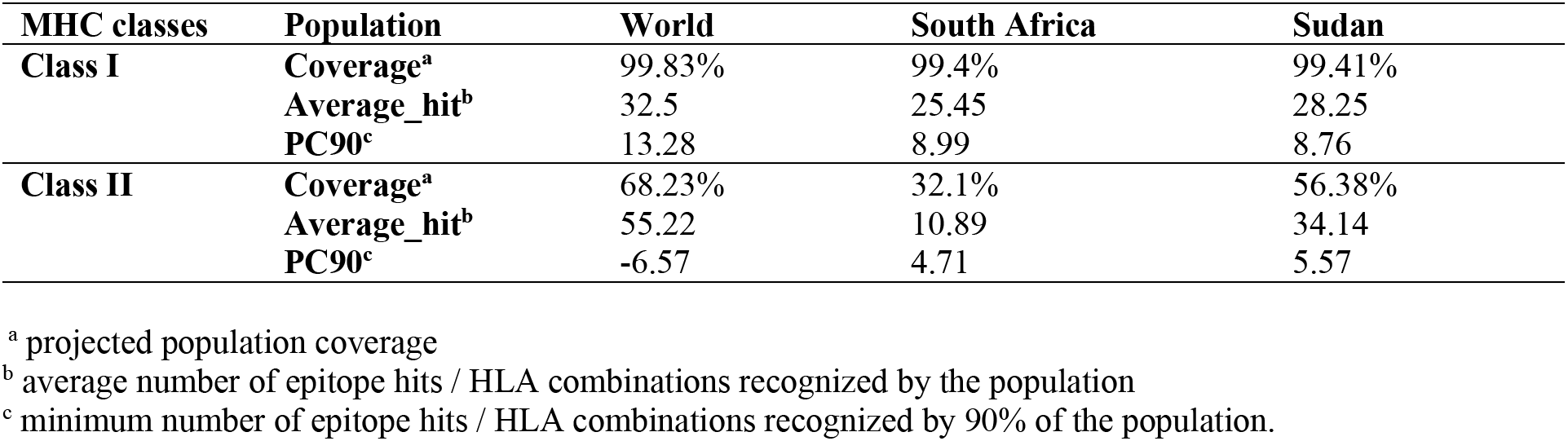
A population coverage for all epitopes that bind to MHC class I and II alleles from different parts of the world.

**Table 5.**
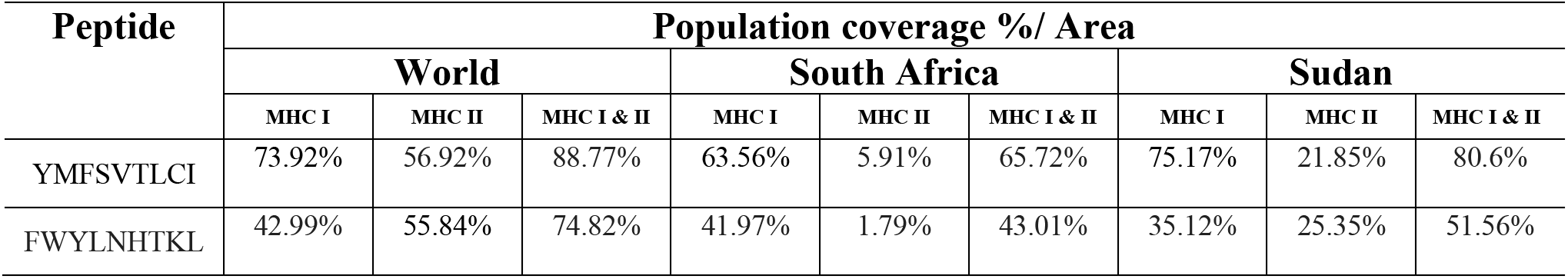
Population coverage of the proposed peptides in MHC class I and MHC class II in five areas

**Figure 3:**
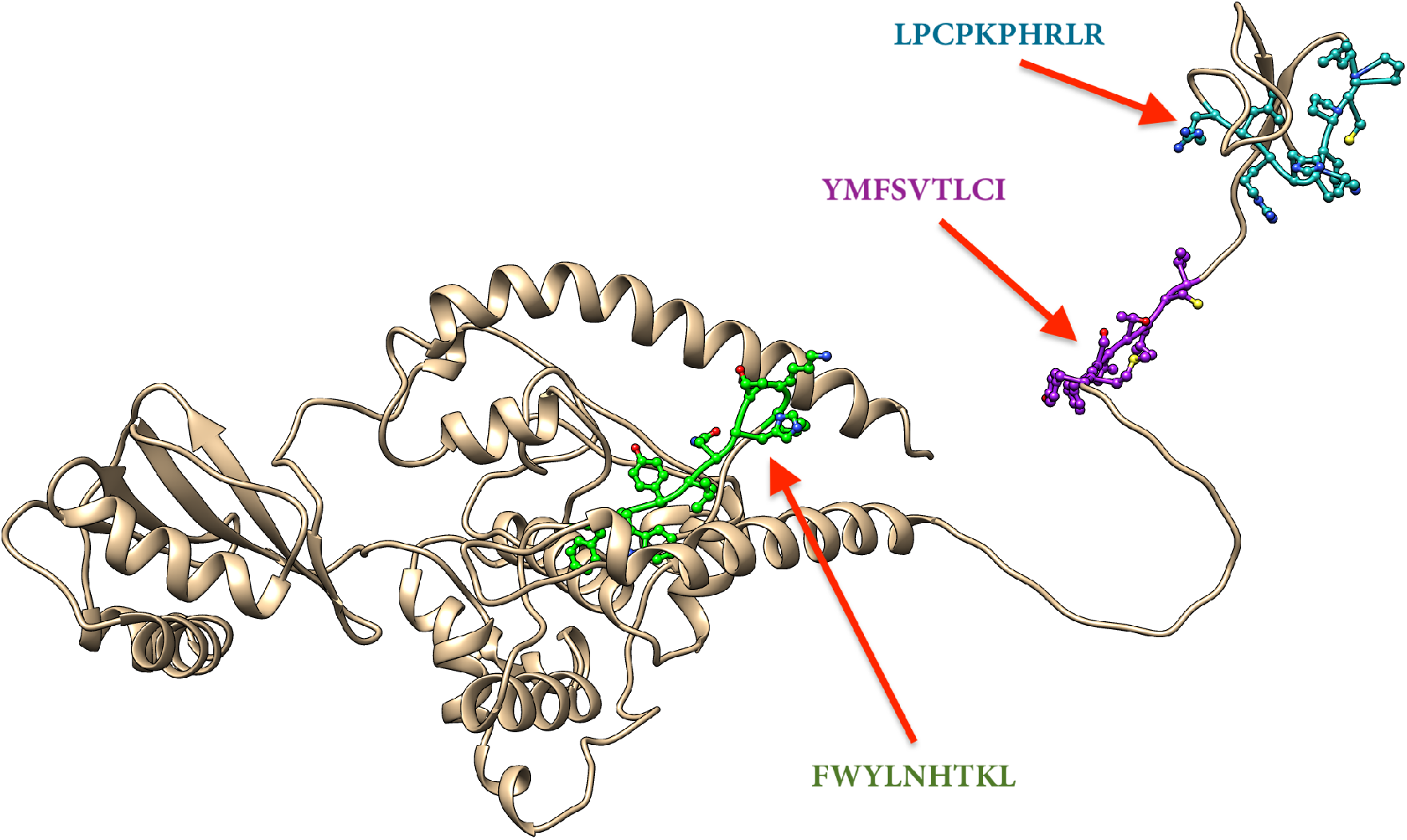
The four potential peptides bound to MHC class I, MHC class II and B cell visualized by Chimera 1.10.2

The binding affinity of the peptide YMFSVTLCI to both class MHC-class I and MHC-class II alleles were found to be 13 different alleles with world population coverage 88.77%, 65.72% for South Africa and 80.6% for Sudan for both MHC class I and II as shown in table 5. According to these interesting findings, a very promising vaccine against *Lujo virus* can be formulated. The most promising three peptides; FWYLNHTKL, LPCPKPHRLR and YMFSVTLCI were docked on to the protein target of GPC domain of *Lujo virus* in complex with the first CUB domain of NRP2 as shown in figure (4 – 6). All peptides were docked on the interface of *Lujo virus* GPC/NRP2 with the following score binding energies: −5.84, −3.88 and −8.20 Kcal/mol for peptides 1, 2 and 3, respectively. Docking results analysis of peptide-1 showed binding hydrogen bonding with two amino acid residues; SER-51 and GLN-131 of NRP2. Peptide-2 showed hydrogen bonding with residues HIS-131 and PHE-137 of *Lujo virus* GPC and ARG-432 of NRP2, while peptide-3 formed hydrogen bonding with PHE-137 of *Lujo virus* GPC. Only Peptide-2 and 3 interacts by forming hydrogen bonding with residues on *Lujo virus* GPC (HIS-131 and PHE-137). These residues are located at the hydrophobic pocket and adjacent to both residues Val139 and Thr140 of α2β4 loop which participates in Van der Waals interactions with NRP2 residues. In addition, histidine residues in the *Lujo virus* GPC/NRP2 complex are obvious candidates for controlling pH-dependent protein–protein interactions. As for peptide-1, it has formed hydrogen bond with GLN-131 of NRP2, which is adjacent to the key residue Arg130 important for NRP2-fc to recognize *Lujo virus* GPC-bearing cells and cell entry of *Lujo virus* ^[2]^. Overall docking results analysis have revealed that the peptides are docked at *Lujo virus* GPC/NRP2 binding surfaces, in which these peptides would serve as potential inhibitors for blocking binding to NRP2 and thus may neutralize the virus. As a result of these interesting outcomes, formulating a vaccine using the suggested peptide is highly promising and encouraging to be highly proposed as a universal epitope-based peptide vaccine against *Lujo virus.*

**Figure 4:**
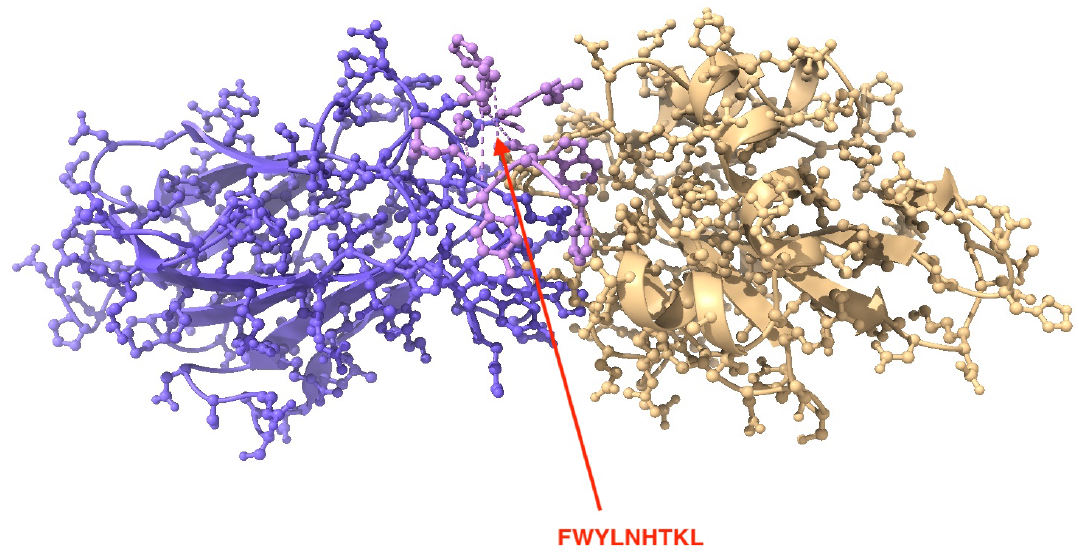
Molecular docking of FWYLNHTKL peptide 1 of *Lujo virus* docked in HLA-A*02:01 and visualized by UCSF Chimera X version 0.1.0.

**Figure 5:**
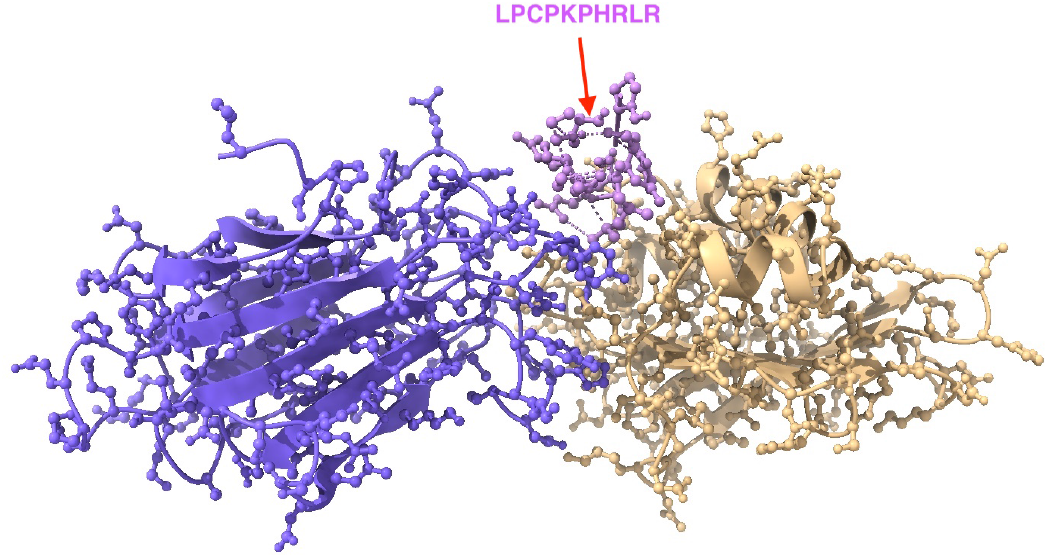
Molecular docking of LPCPKPHRLR peptide 2 of *Lujo virus* docked in HLA-A*02:01 and visualized by UCSF Chimera X version 0.1.0.

**Figure 6:**
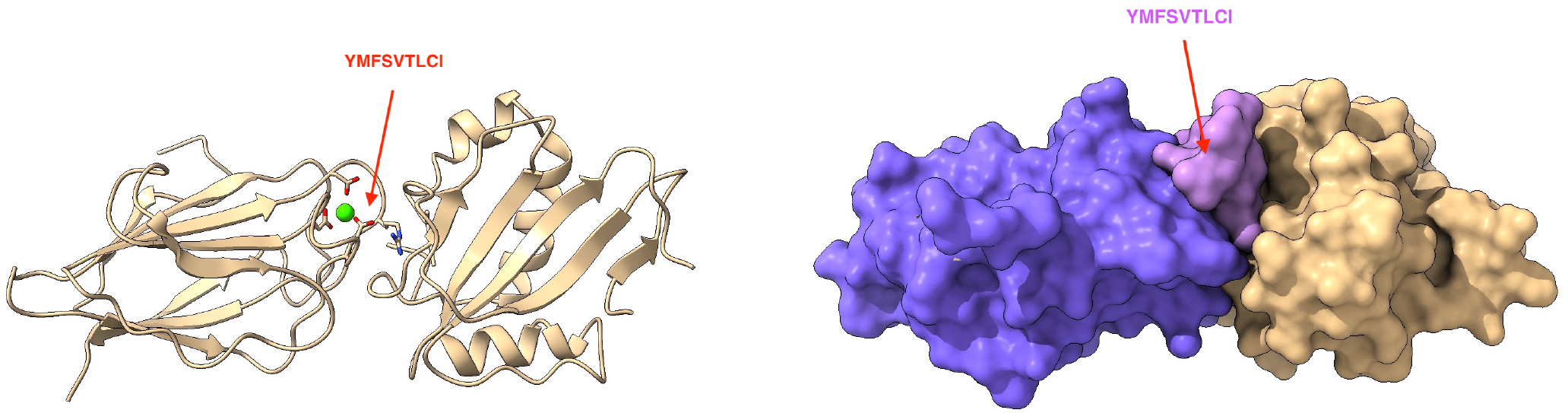
Molecular docking of YMFSVTLCI peptide 3 of *Lujo virus* docked in HLA-A*02:01 and visualized by UCSF Chimera X version 0.1.0 in ribbon and surface structure.

## 5. Conclusions

To the best of our knowledge, this study is considered to be the first to propose an epitope-based peptide vaccine against glycoprotein GPC of *Lujo virus*, which is expected to be highly antigenic with a minimum allergic effect. Furthermore, this study proposes a promising peptide FWYLNHTKL, with a very strong binding affinity to MHC1 and MHC11 alleles. This peptide shows an exceptional population coverage result for both MHC1 and MHC11 alleles.

In-vivo and in-vitro assessment for the most promising peptides namely; FWYLNHTKL, LPCPKPHRLR and YMFSVTLCI are recommended to be explored further to find what ability they may possess to be developed into vaccines against *Lujo virus* glycoprotein GPC.

## References

1. Bergeron, E., et al. (2012). “Reverse genetics recovery of Lujo virus and role of virus RNA secondary structures in efficient virus growth.” J Virol 86(19): 10759–10765.

2. Cohen-Dvashi, H., et al. (2018). “Structural basis for receptor recognition by Lujo virus.” Nat Microbiol 3 (10): 1153–1160.

3. Kunz, S. and J. C. de la Torre (2017). “Breaking the Barrier: Host Cell Invasion by Lujo Virus.” Cell Host Microbe 22 (5): 583–585.

4. Sewlall, N. H., et al. (2014). “Clinical features and patient management of Lujo hemorrhagic fever.” PLoS Negl Trop Dis 8 (11): e3233.

5. Shao, J., et al. (2015). “Human hemorrhagic Fever causing arenaviruses: molecular mechanisms contributing to virus virulence and disease pathogenesis.” Pathogens 4 (2): 283–306.

6. Beyer WR, Popplau D, Garten W, von Laer D, Lenz O. Endoproteolytic processing of the lymphocytic choriomeningitis virus glycoprotein by the subtilase SKI-1/S1P. J Virol (2003);77(5):2866–72. [PubMed: 12584310]

7. Pinschewer DD, Perez M, Sanchez AB, de la Torre JC. Recombinant lymphocytic choriomeningitis virus expressing vesicular stomatitis virus glycoprotein. Proc Natl Acad Sci U S A (2003);100(13): 7895–900. [PubMed: 12808132]

8. Rojek JM, Lee AM, Nguyen N, Spiropoulou CF, Kunz S. Site 1 protease is required for proteolytic processing of the glycoproteins of the South American hemorrhagic fever viruses Junin, Machupo, and Guanarito. J Virol 2008a;82(12):6045–51. [PubMed: 18400865]

9 Tani, H., et al. (2014). “Analysis of Lujo virus cell entry using pseudotype vesicular stomatitis virus.” J Virol88(13): 7317–7330.

12. Mohammed, A. A., et al. (2018). “Epitope-Based Peptide Vaccine Against Fructose-Bisphosphate Aldolase of Madurella mycetomatis Using Immunoinformatics Approaches.” Bioinform Biol Insights12: 1177932218809703.

13. Skwarczynski M, Toth I. Peptide-based synthetic vaccines. Chemical science. (2016);7(2):842–54.

14. Mohammed, A. A., Shantier, S. W., Mustafa, M. I., Osman, H. K., Elmansi, H. E., Osman, I. A. A., Hassan, M. A. (2020). Epitope-Based Peptide Vaccine against Glycoprotein G of Nipah Henipavirus Using Immunoinformatics Approaches. Journal of Immunology Research, 2020. https://doi.org/10.1155/2020/2567957.

15. Badawi MM, Osman MM, Alla AAF, Ahmedani AM, Abdalla Mh, Gasemelseed MM, et al. Highly Conserved Epitopes of ZIKA Envelope Glycoprotein May Act as a Novel Peptide Vaccine with High Coverage: Immunoinformatics Approach. American Journal of Biomedical Research. (2016);4(3):46–60.

16. Li, W., Joshi, M., Singhania, S., Ramsey, K., & Murthy, A. (2014). Peptide Vaccine: Progress and Challenges. Vaccines, 2(3), 515–536.

17. Oyarzun, P., Ellis, J. J., Gonzalez-Galarza, F. F., Jones, A. R., Middleton, D., Boden, M., & Kobe, B. (2015). A bioinformatics tool for epitope-based vaccine design that accounts for human ethnic diversity: Application to emerging infectious diseases. Vaccine, 33(10), 1267–1273.

18. “NCBI protein sequence database,” 2018, https://www.ncbi.nlm.nih.gov/protein.

19. Vita R, Overton JA, Greenbaum JA, Ponomarenko J, Clark JD, Cantrell JR, Wheeler DK, Gabbard JL, Hix D, Sette A, Peters B. The immune epitope database (IEDB) 3.0. Nucleic Acids Res. 2014 Oct 9. pii: gku938. [Epub ahead of print] PubMed PMID: 25300482.

20. Hall TA (1999) BioEdit: a user-friendly biological sequence alignment editor and analysis program for Windows 95/98/NT. Nucl Acids Symp Ser 41: 95–98.

21. Bui, H.H., Sidney, J., Li, W., Fusseder, N., Sette, A., 2007. Development of an epitope conservancy analysis tool to facilitate the design of epitope-based diagnostic sand vaccines. BMC Bioinform. 8 (1), 361.

22. Larsen JE, Lund O, Nielsen M (2006) Improved method for predicting linear B-cell epitopes. Immunome Res 2: 2.

23. Emini, E.A., Hughes, J.V., Perlow, D.S., Boger, J., 1985. Induction of hepatitis A virus-neutralizing antibody by a virus-specific synthetic peptide. J. Virol. 55 (3), 836–839.

24. Kolaskar, A.S., Tongaonkar, P.C., 1990. A semi-empirical method for prediction of anti-genic determinants on protein antigens. FEBS Lett. 276 (1–2), 172–174.

25. Nielsen M, Lundegaard C, Worning P, Lauemøller SL, Lamberth K, et al. (2003) Reliable prediction of T-cell epitopes using neural networks with novel sequence representations. Protein Sci 12: 1007–1017.

26. Kim Y, Ponomarenko J, Zhu Z, Tamang D, Wang P, et al. (2012) Immune epitope database analysis resource. Nucleic Acids Res 40: W525–W530.

27. Zhang Q, Wang P, Kim Y, Haste-Andersen P, Beaver J, et al. (2008) Immune epitope database analysis resource (IEDB-AR). Nucleic Acids Res 36: W513–W518.

28. Källberg M, Wang H, Wang S, Peng J, Wang Z, Lu H, Xu H. Temple-based protein structure modelling using the RaptorX website server. Nature Protocols. (2012); 7: 1511–1522.

29. Goddard TD1, Huang CC, Ferrin. TE Software extensions to UCSF chimera for interactive visualization of large molecular assemblies. J Structure (2005); 13(3):473–82.

30. Choo, J.A.; Thong, S.Y.; Yap, J.; van Esch, W.J.; Raida, M.; Meijers, R.; Lescar, J.; Verhelst, S.H.;Grotenbreg, G.M. Bioorthogonal cleavage and exchange of major histocompatibility complex ligands byemploying azobenzene-containing peptides. Angew. Chem. 2014, 53, 13390–13394.

31. Morris GM, Goodsell DS, Halliday RS, et al (1998) Automated docking using a Lamarckian genetic algorithm and an empirical binding free energy function. J Comput Chem 19:1639–1662. doi: 10.1002/(SICI)1096-987X(19981115)19:14<1639::AID-JCC10>3.0.CO;2-B.

32. Morris GM, Ruth H, Lindstrom W, et al (2009) Software news and updates AutoDock4 and AutoDockTools4: Automated docking with selective receptor flexibility. J Comput Chem 30:2785–2791. doi: 10.1002/jcc.21256.

33. Pettersen EF, Goddard TD, Huang CC, et al (2004) UCSF Chimera - A visualization system for exploratory research and analysis. J Comput Chem 25:1605–1612. doi: 10.1002/jcc.20084.

